# Invasion history reconstruction and potential distribution of the ambrosia beetles *Euwallacea fornicatus* and *E. perbrevis* (Coleoptera: Scolytinae), two global emerging pests

**DOI:** 10.64898/2026.06.01.727051

**Authors:** María Victoria Lantschner, Esteban Ceriani-Nakamurakare, Andrew J. Johnson, Anthony I. Cognato, Sarah M. Smith, Demian F. Gomez

## Abstract

The global spread of invasive insects poses serious ecological and economic threats to forest ecosystems. *Euwallacea fornicatus* and *E. perbrevis* are cryptic ambrosia beetles native to Southeast Asia that have invaded multiple regions worldwide, damaging diverse woody hosts through gallery formation and fungal symbiont inoculation. We compiled confirmed and novel occurrence records to describe their global distributions, reconstruct invasion histories and likely origins using mitochondrial COI phylogenies, and compare their potential distributions through models based on bioclimatic variables. *Euwallacea fornicatus* has expanded rapidly over the past decades, establishing in North America (2003), Israel (2009), South Africa (2016), South America (2020), Australia (2021), Europe (2022) and Turkey (2024). In contrast, *E. perbrevis* has an earlier but slower invasion history, with establishments in Hawaii (1918), Central America (1979), Oceania (1982), and North America (2004). Phylogenetic analyses revealed at least six independent introductions for each species. *Euwallacea fornicatus* primarily originated from native populations in China, Taiwan and Vietnam, whereas *E. perbrevis* from Indonesia and Thailand, with additional introductions from unknown sources. Secondary spread from invaded regions is also likely. Distribution models indicated distinct climatic niches. *Euwallacea fornicatus* tolerates broader thermal ranges and drier conditions, enabling establishment from subtropical to temperate regions, whereas *E. perbrevis* appears restricted to tropical climates. Only 32% of predicted suitable habitat overlapped, indicating low coexistence potential. The broader climatic tolerance and faster recent spread of *E. fornicatus* highlights a higher invasion risk and greater management challenges. These findings provide key insights to strengthen biosecurity strategies aimed at preventing further spread.

**Key message:**

- We updated the global distribution and likely invasion routes of two cryptic ambrosia beetles.
- We identified multiple independent and secondary introductions worldwide.
- We revealed distinct climatic niches and limited habitat overlap.
- We showed higher invasion risk for *E. fornicatus* due to broader climatic tolerance.
- We identified high-risk regions for surveillance based on climate suitability.

## Introduction

The expansion of global trade has significantly increased the frequency of biological invasions by forest insects in recent decades, as many species are unintentionally transported through wood products, live plants, and packaging materials (Seebens et al. 2017; Brockerhoff and Liebhold 2017; Meurisse et al. 2019). Once established, invasive forest insects can cause profound ecological impacts—altering forest composition, reducing biodiversity, and disrupting key ecosystem functions—as well as substantial economic losses related to timber production, forest health management, and the provision of ecosystem services (Aukema et al. 2011; Boyd et al. 2013). Understanding the invasion history of these species and accurately predicting their potential distribution in new regions are critical steps for anticipating future introductions and the spread of potential pests (Hulme 2009; Venette et al. 2010). This knowledge is essential for informing surveillance programs, prioritizing quarantine efforts, and implementing proactive measures to prevent invasions (Early et al. 2016; Seebens et al. 2018).

Among the most concerning invasive forest insects are ambrosia beetles (Coleoptera: Curculionidae: Scolytinae and Platypodinae), several of which have emerged as significant pests due to their unique ecological strategy and their devastating impact on both forest and agricultural systems. Ambrosia beetles cultivate symbiotic fungi within host trees as their primary food source. This mutualistic relationship allows even small beetle populations to cause disproportionate ecological and economic harm, especially in naive environments lacking coevolved defenses (Hulcr and Stelinski 2017; Mech et al. 2019). Their broad host range — including both native and non-native, as well as ornamental or timber production species—facilitates their establishment across diverse ecosystems, while their cryptic morphology and hidden wood-boring habits make detection and management particularly challenging (Lantschner et al. 2020; Grégoire et al. 2023). The recent global spread of several ambrosia beetle species, such as *Xylosandrus crassiusculus, Xylosandrus germanus, Euwallacea interjectus*, and *Euwallacea fornicatus*, highlights their capacity to rapidly invade new biogeographic regions, exploiting novel hosts, and demonstrating an alarming ability to expand its ecological niche (Cognato et al. 2015; Dzurenko et al. 2021; Urvois et al. 2023; Lai et al. 2024; Ceriani-Nakamurakare et al. 2025).

*Euwallacea* includes a complex of ambrosia beetles known for their invasive potential, primarily due to their mutualistic associations with fungal symbionts that may cause damage to a wide range of woody plant hosts (Eskalen et al. 2013; O’Donnell et al. 2015). Two of the most ecologically and economically important members of this group are *E. fornicatus* (polyphagous shot hole borer, PSHB) and *E. perbrevis* (tea shot hole borer, TSHB). These cryptic species have expanded well beyond their native range in Southeast Asia over recent decades, largely through human-mediated transport of infested wood and plant materials (Gomez et al. 2018; Smith et al. 2019; Ceriani-Nakamurakare et al. 2023).

Genetic studies have revealed that both species have undergone multiple introduction events involving distinct source populations (Stouthamer et al. 2017; Covre et al. 2024; Ceriani-Nakamurakare et al. 2025). This complex invasion history is likely driven by international trade and the global movement of wood products. Although these species are known to share morphological similarities and overlapping host ranges, they appear to differ in their ecological tolerances. Determining whether they occupy different climatic niches is essential for improving invasion forecasts and strengthening risk assessments. Climatic variables such as temperature and humidity usually play a significant role in limiting or facilitating species dispersal and may help explain divergent invasion dynamics, even among closely related taxa. The objectives of this study are: (1) to describe current distributions of *E. fornicatus* and *E. perbrevis* at a global scale, reconstruct their invasion history and identify their points of origin worldwide based on an analysis of genetic structure; and to (2) describe and compare the potential distribution of both species based on bioclimatic variables.

## Materials & Methods

### Current distribution

We compiled a database of occurrence locations of *E. fornicatus* and *E. perbrevis* across their native and non-native range from published papers and unpublished reports from experts. It is acknowledged that there are difficulties in identifying these species based solely on morphological characteristics. For this reason, we have included only genetically confirmed records or samples that have been identified by specialists whose morphological identifications have been previously validated through molecular data (see Table S1 for details). To characterize the invasion chronology beyond the native range, we compiled the year of first detection for each species and recorded whether it was eradicated or subsequently confirmed as established (i.e. species recorded over consecutive years and at multiple locations within the invaded region) in each non-native country.

### Molecular phylogenetics and global haplotype distribution

From the compiled species occurrence data, we selected those records of *E. fornicatus* and *E. perbrevis* for which mitochondrial gene cytochrome oxidase subunit I (COI) sequences were available (Table S1). For newly sequenced specimens, DNA extraction and COI amplification were performed using primers COI_1455b/COI_r750 according to Smith and Cognato (2014). Amplified products were Sanger sequenced and sequences were assembled in Sequencher (Ann Arbor, MI). Illumina sequencing was also used to generate sequences for older or degraded specimens (Cognato et al. unpublished). All sequences were deposited in GenBank under accession numbers: PX978039-PX978043, PX977808-PX977811. All phylogeographic analyses were conducted at the country level including their insular territories, except American Samoa (ASM) that was treated independently because it is not recognised as a state of the United States of America (USA). Country codes follow the ISO 3166-1 alpha-3 standard.

Haplotype network reconstruction employed mitochondrial COI sequences that were aligned using MUSCLE v5 (Edgar 2022). For specimens with available genome assemblies (e.g*., E. fornicatus* EFF26), the COI gene was extracted from the mitochondrial genome with default parameters via the R package ‘orthGS’ (Aledo and Aledo 2025)and aligned with the COI database as described above. The final alignments comprised 154 sequences (569 bp) for *E. fornicatus* and 64 sequences (569 bp) for *E. perbrevis*; gaps were excluded from all the analyses. Haplotype identification and genetic differentiation estimates (e.g. pairwise FST, GST, nucleotide diversity, among others) were calculated in DnaSP v6.12 (Rozas et al. 2017). A new haplotype numbering system has been established for this study, and the complete results are provided in Tables S3–S6. Statistical parsimony haplotype networks were constructed using the TCS method (Clement et al. 2000) implemented in PopART (Leigh et al. 2015) to visualize genealogical relationships among haplotypes. When referring to haplotype designations originally described by Stouthamer et al. (2017), Wang et al. (2022), and other curated sources, former identifiers are provided in parentheses (e.g. ‘formerly H33’).

Maximum-likelihood phylogenetic trees (ML) were constructed in IQ-TREE v2.0 (Minh et al. 2020) using 1000 standard bootstrap replicates and 1000 SH-like approximate likelihood-ratio test (SH-aLRT) replicates (Anisimova and Gascuel 2006). Each species dataset included a single representative sequence per haplotype, with *E. fornicatior* (MH276931) used as an outgroup. Best-fit substitution models were determined using ModelFinder (Kalyaanamoorthy et al. 2017) according to the Bayesian Information Criterion value, yielding TPM2u+F+I+G4 for *E. fornicatus* and TIM2+F+R3 for *E. perbrevis*.

A normalized heatmap was constructed by dividing each haplotype’s distribution equally among all countries where it occurred (row sum= 100%), to emphasize distribution breadth independent of sequencing and/or sampling intensity. Geographic occurrences and relative frequencies of the identified mitochondrial haplotypes for *E. fornicatus* and *E. perbrevis* were mapped globally. Haplotype proportions within each country were depicted using pie charts to illustrate the spatial partitioning of genetic diversity.

### Potential distribution

Species distribution models were developed for *E. fornicatus* and *E. perbrevis* using occurrence records compiled across both their native and non-native ranges. Only records representing confirmed established populations were retained, and greenhouses occurrences were excluded. To reduce spatial bias and improve the uniformity of occurrence data, occurrences were spatially thinned using aa 5-minute distance radius, resulting in 98 presence records for *E. fornicatus* and 53 presence records for *E. perbrevis*. We considered climate conditions as potential predictors of the distribution of both species, based on 19 bioclimatic variables from WorldClim v2.1, spanning 1970 to 2000, at 5 arc-min resolution (Fick and Hijmans 2017). In order to avoid collinearity among the variables, a matrix of Spearman’s rank correlation coefficients for all possible pairs of variables was carried out for each occurrence location of the species using the R package Hmisc (Harrell Jr 2026). Variables that correlated (r≥ 0.7) with each other were excluded, leaving only those variables with more biological relevance for the species. The selected variables were: annual mean temperature (BIO1), mean diurnal range (BIO2), isothermality (BIO3), annual precipitation (BIO12), precipitation seasonality (coefficient of variation, BIO15).

A common background extent was defined for both species to ensure comparability. A binary mask was constructed encompassing the native biogeographic region (Indo-Malayan) and invaded areas, including a 100 km buffer around occurrences. A total of 1,000 background points were randomly sampled within this area for each species (Barve et al. 2011). Models were fitted independently for each species using the biomod2 package. Four algorithms were implemented: Generalized Linear Models (GLM), Random Forest (RF), Gradient Boosting Machines (GBM), and Maximum Entropy (MaxEnt). Models were fitted independently for each species. GLMs used a binomial error distribution with AIC-based stepwise selection. RF models included 1,000 trees, with √p variables per split and a minimum node size of 5. GBMs were fitted with 2,000 trees, interaction depth = 3, learning rate = 0.01, and bag fraction = 0.75, with internal cross-validation to optimize iteration number. MaxEnt complexity was tuned separately for each species using a grid search across regularization multipliers (0.5–5) and feature class combinations (linear, quadratic, hinge, product). The optimal configuration was selected based on the highest mean AUC under 5-fold cross-validation.

Model performance was evaluated using 5-fold cross-validation with 5 repetitions (25 replicates per algorithm). Model skill was assessed using AUC and TSS, and only models with TSS ≥ 0.4 were retained. Ensemble models were constructed using a TSS-weighted mean, where each replicate contributed proportionally to its TSS score. Variable importance was estimated using permutation tests, averaging the decrease in AUC across permutations, replicates, and algorithms, and rescaling values to sum to 100%. Ensemble models were projected globally to identify climatically suitable areas. Final suitability maps represent TSS-weighted mean predictions scaled from 0 to 1. To estimate the potential geographical distribution of *E. fornicatus* and *E. perbrevis* outside of its native range, we designated a threshold probability to define suitable and non-suitable habitat using TSS-maximizing thresholds. Marginal response curves were generated for all predictors using biomod2, holding non-focal variables constant at their mean values. Curves represent the mean prediction across models, with standard deviation indicating inter-model uncertainty.

Climatic niche overlap between *E. fornicatus* and *E. perbrevis* was quantified in both geographic and environmental space. In geographic space, Schoener’s D was calculated from normalized ensemble suitability maps after aligning raster resolution and extent. In environmental space, overlap was assessed using a PCA of environmental variables based on a random sample of 10,000 background points. Occurrence records for both species were projected onto the first two principal components, and species densities were estimated using a kernel density approach. Overlap metrics were then computed from these density grids. Niche equivalency and similarity tests were performed using 100 random permutations to evaluate whether niches were identical and whether observed overlap differed from random expectations.

All analyses were conducted in R, using *biomod2* (Guéguen et al. 2026) for distribution modeling, *maxnet* (Phillips et al. 2017) for MaxEnt tuning, *terra* (Hijmans et al. 2026) for raster processing, *sf* (Pebesma 2018) for vector data, and *ecospat* (Di Cola et al. 2017) for niche overlapping. Additional spatial processing and map layouts were performed in ArcGIS Pro 3.6 (Esri, Redlands, CA, USA).

## Results

### Current distribution

We found confirmed records of *E. fornicatus* within its native range across Southeast Asia, including China, India, Japan, Laos, Malaysia, Sri Lanka, Taiwan, Thailand, and Vietnam (Fig. 1, Table S1). The earliest confirmed establishment outside the native range occurred in the United States of America, with detections in California (2003) and Hawaii (2006). Subsequent invasions were reported in Israel (2009), South Africa (2016), Brazil (2020), Argentina and Australia (2021), Spain (2022), Uruguay (2023), and Turkey (2024) (Fig. 1, Table S1). Additional records from Poland (2017), Italy (2020), and Germany and the Netherlands (2021) corresponded to confined occurrences in greenhouses or botanical gardens that were successfully eradicated.

**Fig 1.**
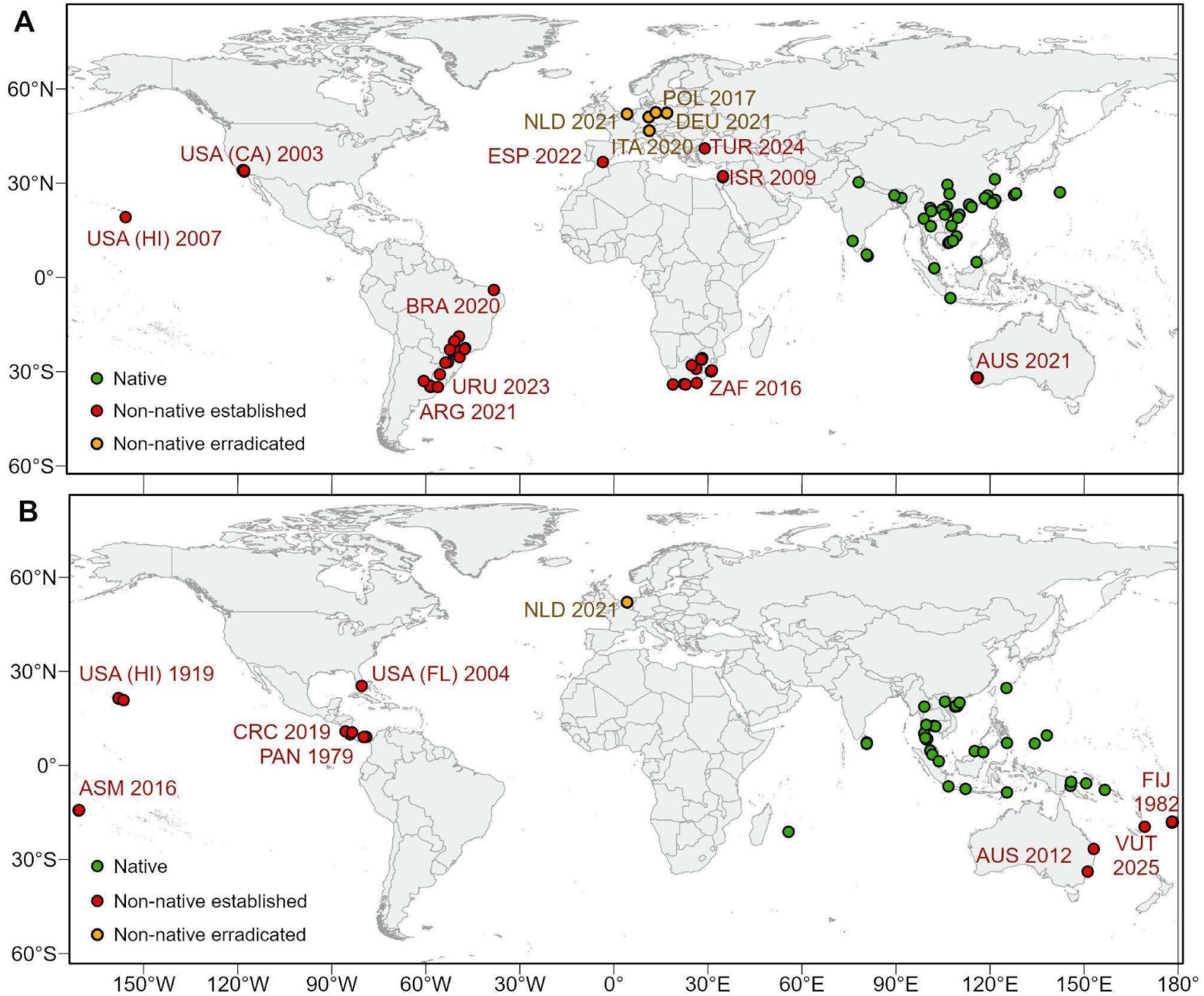
Current global distribution of (A) *Euwallacea fornicatus* and (B) *E. perbrevis*. Green circles indicate records within the native range, red circles indicate non-native established populations, and orange circles indicate non-native eradicated populations. The year of first detection is shown for each non-native country

For *E. perbrevis*, we found confirmed records within its native range across Southeast Asia, including Brunei, China, Indonesia, Japan, Malaysia, the Philippines, Singapore, Sri Lanka, Thailand, Timor-Leste, and Vietnam; several islands in Oceania, including the Federated States of Micronesia, Palau, Papua New Guinea, and the Solomon Islands; and Réunion Island in the Indian Ocean (Fig. 1, Table S1). Although the precise native range of *E. perbrevis* remains uncertain and some island populations may have experienced early human-mediated dispersal, all were treated here as part of the native range due to geographical proximity. The earliest non-native records occurred in Oceania, including Hawaii (1918), Fiji (1982), eastern Australia (2012), American Samoa (2016) and Vanuatu (2025). In the Americas, the species was first detected in Panama (1979), followed by the United States of America (Florida, 2004) and Costa Rica (2019) (Fig. 1, Table S1). A single introduction in the Netherlands (2021) was detected and subsequently eradicated.

### Molecular phylogenetics and global haplotype structure

Maximum likelihood phylogenetic reconstruction of *E. fornicatus* and *E. perbrevis* species revealed distinct biogeographic structuring, with varying degrees of nodal support across the tree topology. Sequences from putative native ranges exhibited extensive paraphyly, occupying basal and internal nodes throughout the phylogenies (Figs. 2, 3 and S1).

**Fig 2.**
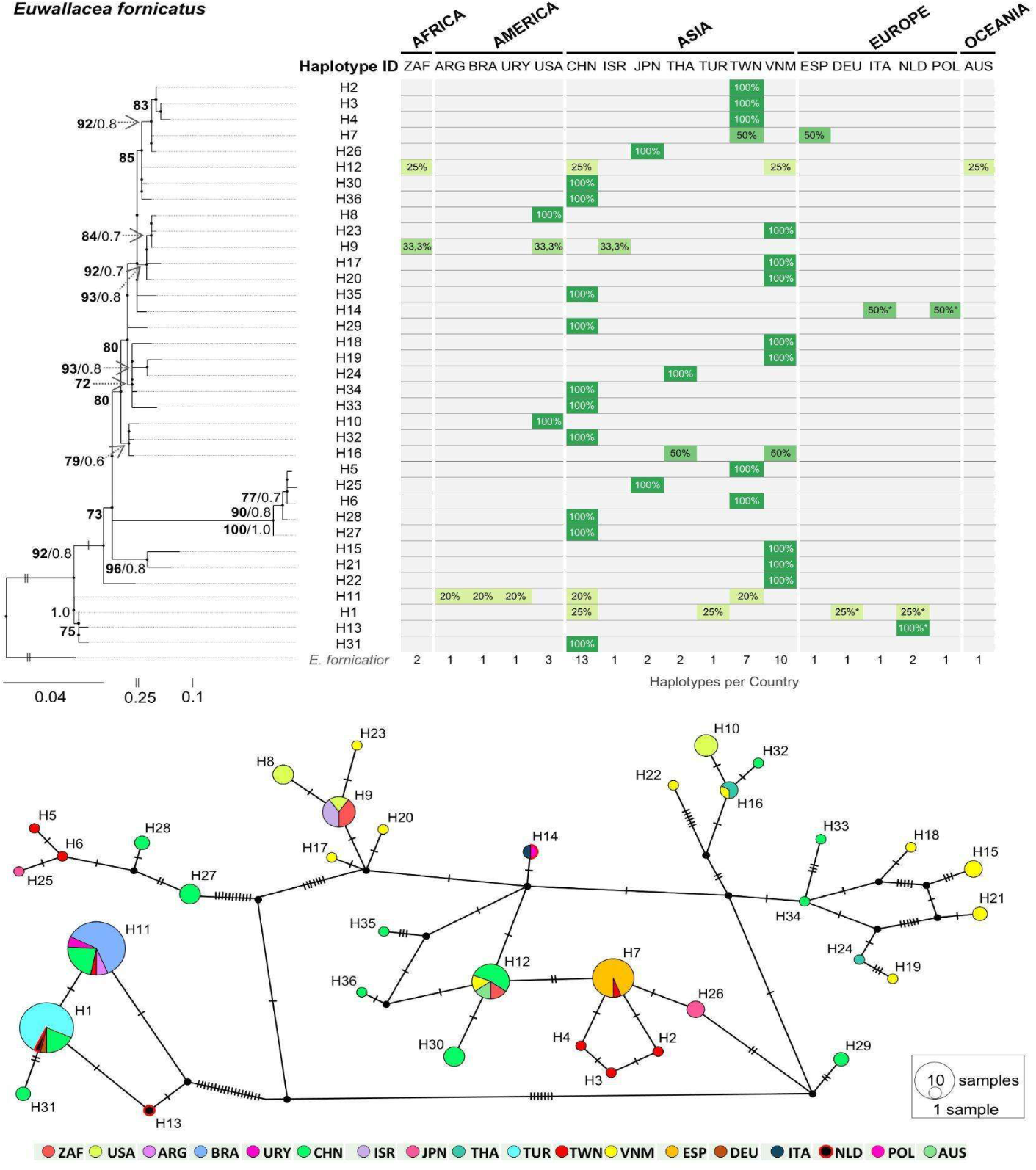
Phylogeographic structure and haplotype distribution of *Euwallacea fornicatus*. (Top) Maximum likelihood phylogenetic tree of 36 haplotypes (H1–H36), with *E. fornicatior* as outgroup. Node support values denote Bootstrap and SH-aLRT probabilities (>60% in bold and >0.5, respectively). Branch lengths represent nucleotide substitutions. The accompanying heatmap shows the proportional occurrence (%) of each haplotype across surveyed countries; cells with 100% indicate haplotypes exclusive to a given country. (Bottom) TCS haplotype network, where node size is proportional to sample frequency and colors denote countries of origin. Cross-hatches along connecting branches represent mutational steps between haplotypes. Asterisks indicate eradicated populations. Country codes: ARG: Argentina, AUS: Australia, BRA: Brazil, CHN: China, DEU: Germany, ESP: Spain, ISR: Israel, ITA: Italy, JPN: Japan, NLD: Netherlands, POL: Poland, THA: Thailand, TUR: Turkey, TWN: Taiwan, URY: Uruguay, USA: United States of America, VNM: Vietnam, ZAF: South Africa

For *E. fornicatus*, we identified 36 distinct haplotypes (H1 to H36, Fig. 2, Table S2), with high global haplotype diversity (Hd= 0.85), and significant genetic differentiation among populations (FST= 0.7; P< 0.001). Both maximum likelihood phylogenetic reconstruction and TCS haplotype network suggest China and Vietnam as the primary reservoirs of genetic diversity (Figures 2 and S1). Asian populations exhibited marked polyphyly. In contrast, non-native populations in South America (Argentina, Brazil, and Uruguay) were strictly monomorphic, all sharing haplotype H11, which was also present in China and Taiwan. Pairwise comparisons among South American countries showed no genetic differentiation (FST= 0, Dxy= 0). South Africa showed a heterogeneous haplotype composition, with co-occurrence of haplotypes H9 and H12, while Israel was monomorphic for H9. These populations clustered together with continental United States of America (USA) within a supported clade (Bootstrap/SH-arlt: 84/0.7). On the other hand, haplotype H12 in was also shared with China, Vietnam, and Australia (Fig. 2).

In the USA, we found strong genetic differentiation between continental USA and Hawaiian populations. Mainland USA displayed haplotypes H8 and H9, whereas Hawaii was monomorphic for H10. Pairwise comparisons indicated near-complete genetic differentiation between the two regions (FST= 0.9, Dxy= 3, Da= 0.02, Table S3). Continental populations also exhibited higher haplotype diversity (Hd= 0.385) than Hawaii. Haplotype H7 was shared within Spain and Taiwan, with a well-supported lineage (Bootstrap/SH-arlt: 77/0.7). Haplotype H1 was shared within China, the Netherlands, Germany, and Turkey, while haplotype H14 was shared within Italy, and Poland which clustered with high support (Bootstrap/SH-arlt: 93/0.8) at distal nodes. These lineages showed strong genetic differentiation when compared to the South American clade (FST≈ 0.8, Dxy= 0.05, Table S3).

For *E. perbrevis* we identified a total of 27 haplotypes (Fig. 3, Table S4), with high global haplotype diversity (Hd= 0.9) and significant differentiation among populations (FST= 0.7. P< 0.001, Table S5). The ML reconstruction identified two primary clades, which were primarily rooted in the genetic diversity of Thailand and Indonesia, which together harboured the higher haplotype richness, with 10 and 4 unique haplotypes, respectively (Fig. 3). These clades showed strong genetic differentiation (FST≈ 0.8). Papua New Guinea exhibited haplotypes that clustered separately with both Indonesian and Thai lineages and exhibited high intrapopulation divergence (Hd= 1, π= 0.1), with haplotypes H9 and H12 separated by 56 mutational steps.

**Fig 3.**
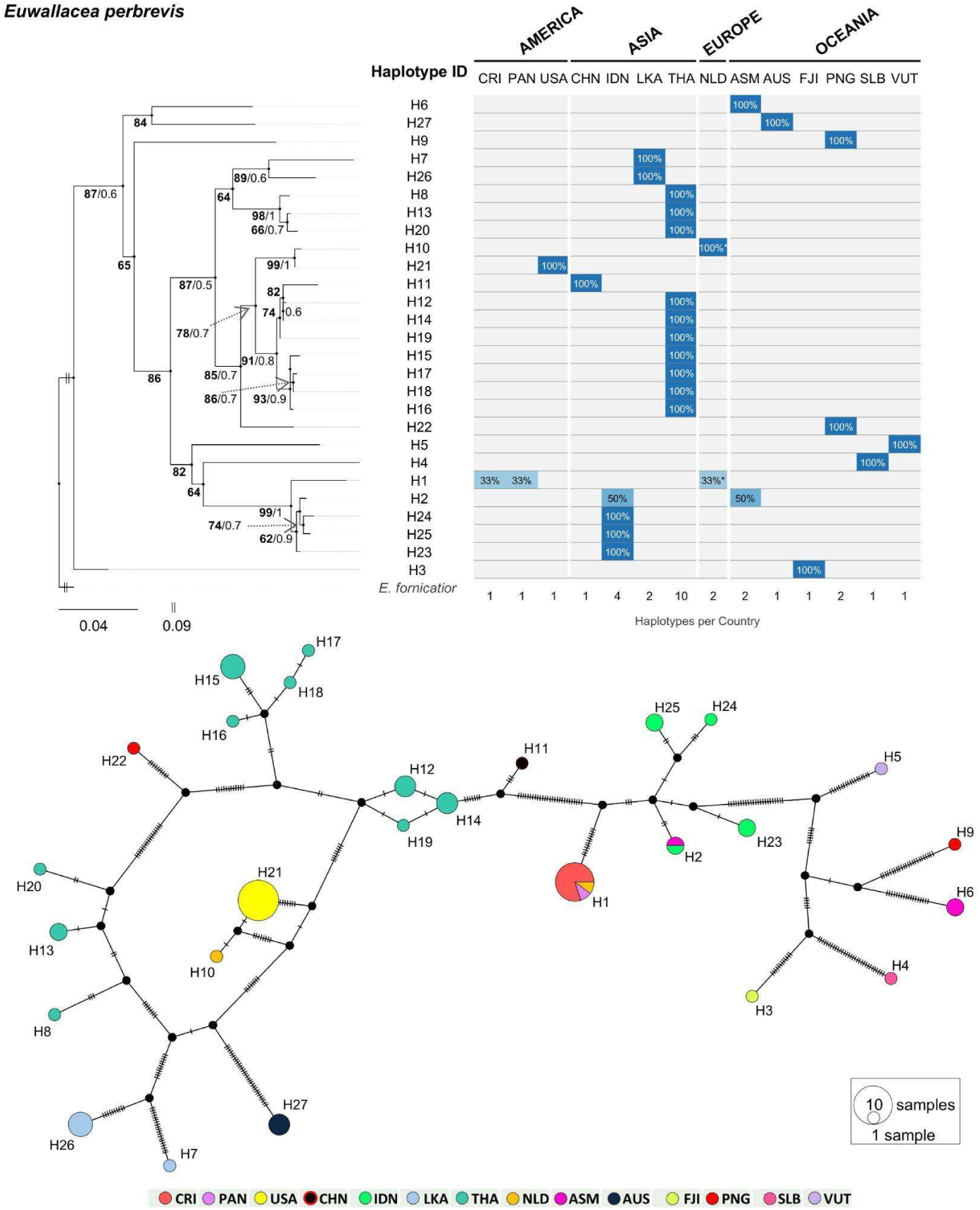
Phylogeographic structure and haplotype distribution of *Euwallacea perbrevis*. (Top) Maximum likelihood phylogenetic tree of 27 haplotypes (H1–H27), with *E. fornicatior* as outgroup. Node support values denote Bootstrap and SH-aLRT probabilities (>60% in bold and >0.5, respectively). Branch lengths represent nucleotide substitutions. The accompanying heatmap shows the proportional occurrence (%) of each haplotype across surveyed countries; cells with 100% indicate haplotypes exclusive to a given country. (Bottom) TCS haplotype network, with node size proportional to sample frequency and colors denote countries of origin. Cross-hatches across connecting branches represent mutational steps between haplotypes. The asterisk indicates an eradicated population. Country codes: AUS: Australia, ASM: American Samoa, CHN: China, CRI: Costa Rica, FJI: Fiji, IDN: Indonesia, LKA: Sri Lanka, NLD: Netherlands, PAN: Panama, PNG: Papua New Guinea, SLB: Solomon Islands, THA: Thailand, USA: United States of America, VUT: Vanuatu

Conversely, nine populations were monomorphic: Australia, China, Costa Rica, Fiji, Panama, Solomon Islands, USA, Vanuatu, and the eradicated population from the Netherlands. Among these, Costa Rica, Panama and the Netherlands shared haplotype H1 and showed no genetic differentiation (FST= 0 across all pairwise comparisons, Table S5). The Netherlands population was genetically identical to the USA (FST= 0, Bootstrap/SH-aLRT: 99/1). In contrast, the USA haplotype exhibited complete differentiation from both the Central American and the Chinese populations (FST= 1, Bootstrap/SH-aLRT: 99/1 and 78/0.7, respectively; Dxy= 0.05), suggesting significant divergence despite their placement within the same major phylogenetic clade.

### Potential invasion routes

Based on the distribution of haplotypes, phylogenetic relationships and dates of first detection in each region, we proposed a set of plausible invasion routes for both species (Fig. 4). For *E. fornicatus*, eight haplotypes were identified in non-native regions. The earliest invasion occurred in California (USA) in 2003, involving haplotypes H9 and H8 (formerly H33 and H35). H9 likely originated from Vietnam, while H8 (formerly H35) has not been detected in the native range but is closely related to H9 (Fig. 2 and 4A). H9 subsequently appeared in Israel and later in South Africa, suggesting introductions either directly from its native range or from previously established populations. Hawaii was colonized by haplotype H10 (formerly H43), which is absent from sampled native populations but closely related to haplotypes from Vietnam, Thailand, and China (Yunan; Fig. 2). South Africa experienced a second introduction of haplotype H12 (formerly H38), likely from Vietnam and/or China (Hong Kong). This same haplotype was later reported in Australia (2021), indicating either a direct introduction from the native range or secondary dispersal from South Africa. In South America, all populations shared haplotype H11 (formerly H52), likely originated from China (Hainan and Fujian) and/or Taiwan. In the Netherlands and Germany haplotype H1 was likely introduced from China (Hainan) and subsequently eradicated. However, the same haplotype was later found to be established in Turkey in 2024, and consequently these introductions could have been interconnected. In Italy and Poland introduction involved haplotype H14, which was not sampled from the native range and was later eradicated. In Spain haplotype H7 (formerly H39) was introduced in 2022, most likely originated from Taiwan.

**Fig 4.**
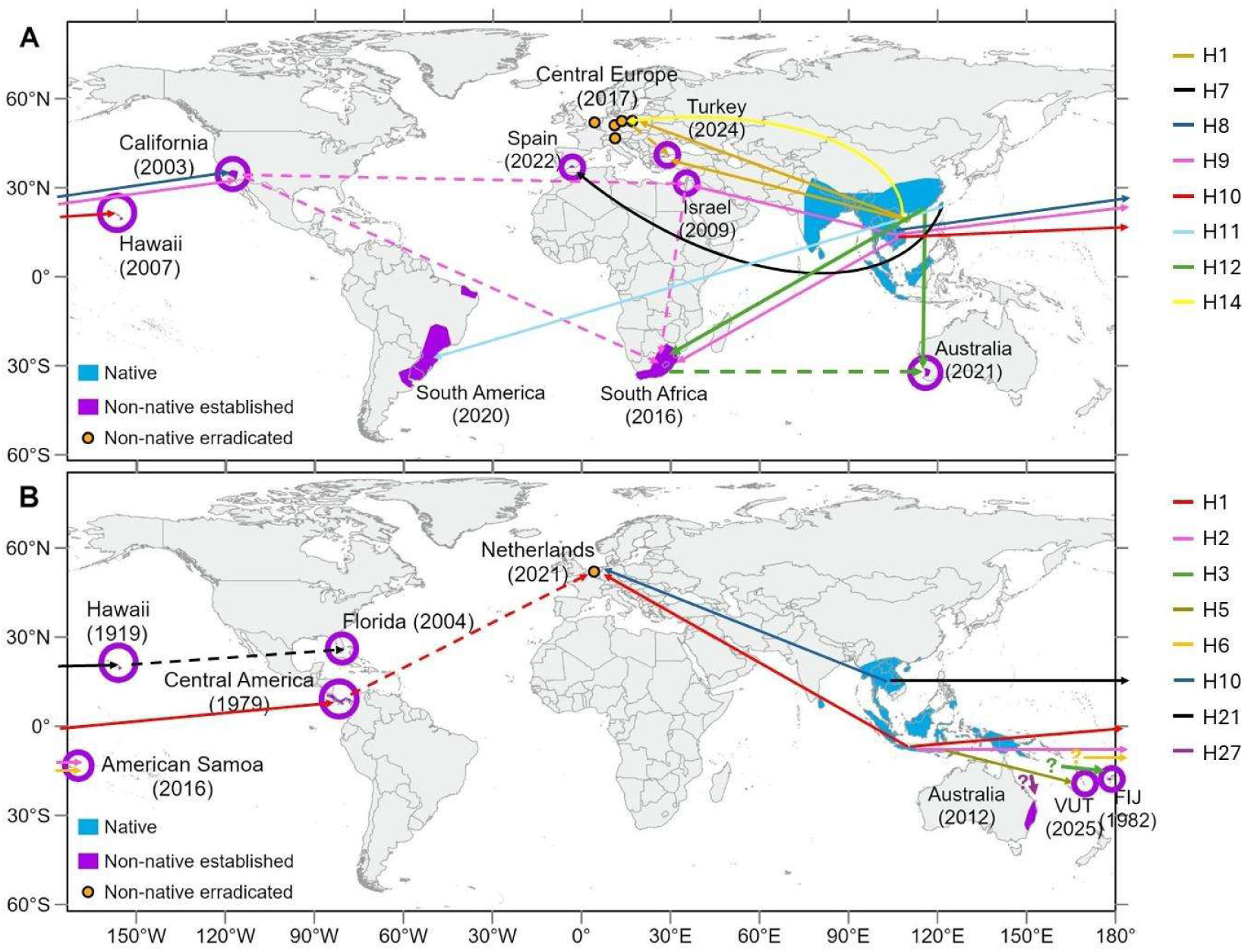
Global reconstruction of the most probable invasion routes of *Euwallacea fornicatus* (A) and *E. perbrevis* (B), inferred from haplotype distributions across native and invaded regions and the temporal sequence of invasion events. Colored lines indicate hypothesized invasion routes associated with specific haplotypes (see Figures 2 and 3). Years in parentheses denote the first reported occurrence in each region outside the native range

For *E. perbrevis*, eight haplotypes were detected in non-native regions. The earliest invasion lineage was detected in Hawaii in 1919 involving haplotype H21, likely originating from Thailand (Fig 3 and 4B). This haplotype was later recorded in the continental United States of America (Florida), likely introduced from secondary populations from Hawaii or directly from the native populations. More recently a phylogenetically closely related haplotype (H10), probably also originated from Thailand, was introduced to the Netherlands and subsequently eradicated. Haplotype H1, detected in Panama and subsequently detected in Costa Rica and the Netherlands (eradicated), as well as haplotype H5 detected in Vanuatu, are associated with a second lineage likely native to Indonesia (Java), The distribution of H1 across these regions may represent successive introductions from already invaded populations. In American Samoa haplotype H2 was likely also introduced from Indonesia, while H6 was introduced from another unknown source, so far restricted to this location. The haplotypes introduced to southeastern Australia (H27) and Fiji (H3) belonged to distinct lineages, with no clear source population identified (Fig. 4).

### Potential distribution

All models exceeded the minimum performance thresholds (AUC> 0.7; TSS> 0.4) and were retained for ensemble construction. Calibration performance was consistently high for both species, with AUC ≥ 0.930 and TSS ≥ 0.727 for *Euwallacea fornicatus*, and AUC ≥ 0.959 and TSS ≥ 0.831 for *E. perbrevis* (Table S6). MaxEnt parameter tuning identified different optimal configurations for each species: RM = 1 with FC = LQHP for *E. fornicatus*, and RM = 0.5 with FC = LQ for *E. perbrevis* (Table S7).

TSS-based weights were similar across algorithms in both species, with RF contributing the highest proportion (29.6% in *E. fornicatus*; 27.4% in *E. perbrevis*), followed by GBM, GLM, and MaxEnt with comparable contributions (Table S8). Ensemble models showed high predictive performance. For *E. fornicatus*, the weighted mean ensemble achieved AUC = 0.953, TSS = 0.781, and Boyce Index = 0.988; for *E. perbrevis*, AUC = 0.969, TSS = 0.842, and Boyce Index = 0.949.

At the global scale, both species showed broad areas of climatically suitable habitat, although with different spatial patterns (Fig. 5A–B). *Euwallacea perbrevis* was mainly associated with tropical regions, whereas *E. fornicatus* extended into subtropical and temperate areas. The predicted suitable area also differed between species (Fig. 5C): *E. fornicatus* occupied 64% of the total area identified as suitable for either species, while E. *perbrevis* occupied 68%. Only 32% of this area was suitable for both species simultaneously.

**Fig 5.**
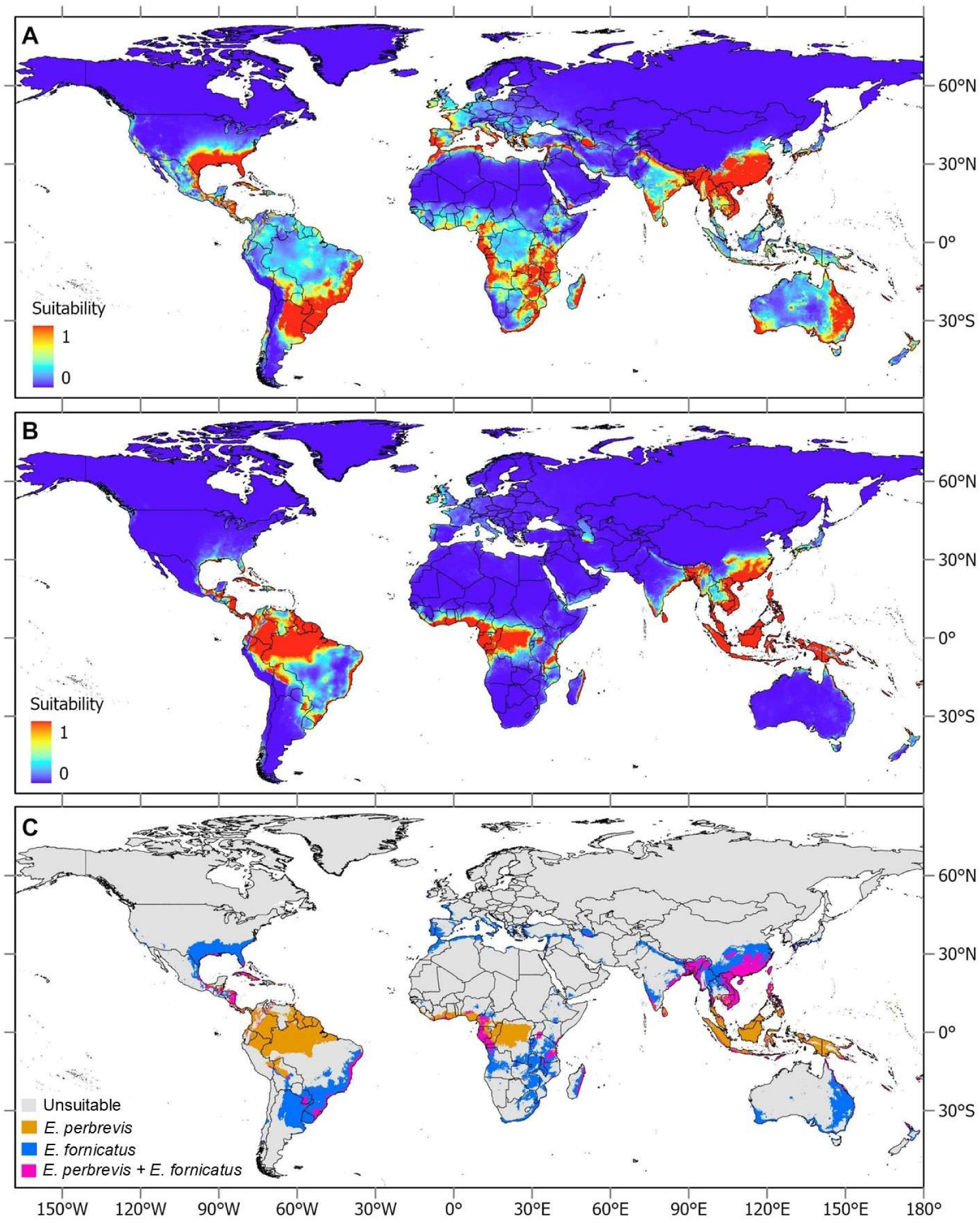
Predicted distribution of the studied species, based on the TSS-weighted ensemble models (GBM, GLM, MaxEnt, and RF). (A) Predicted habitat suitability for *Euwallacea fornicatus.* (B) Predicted habitat suitability for *Euwallacea perbrevis*. (C) Combined predicted distributions of *E. fornicatus* and *E. perbrevis*. Orange polygons indicate areas where only *E. perbrevis* is predicted to find suitable conditions, blue polygons represent areas suitable only for *E. fornicatus*, and purple polygons highlight regions where both species are predicted to be able to coexist

Mean annual temperature (BIO1) was the most important predictor for both species, followed by mean diurnal temperature range (BIO2), while the remaining variables showed lower contributions (Table 1). Response curves indicated a unimodal relationship between temperature and suitability for *E. fornicatus*, with a peak at approximately 18–22 °C. In contrast, *E. perbrevis* showed increasing suitability toward higher temperatures (>20–25 °C) without a clear decline at the upper range. The suitable range of mean diurnal temperature varied from 0–16 °C for *E. fornicatus* and from 0–11 °C for *E. perbrevis* (Fig. S2).

**Table 1.** Mean variable importance scores based on permutation analysis, averaged across all models and cross-validation replicates for *E. fornicatus* and *E. perbrevis*.

| Variable | *E. fornicatus* | *E. perbrevis* |
| --- | --- | --- |
| Annual mean temperature | 55.9 | 37.6 |
| Mean diurnal range | 17.2 | 29 |
| Isothermality | 14.4 | 13.4 |
| Annual precipitation | 9.2 | 13.3 |
| Precipitation seasonality | 3.2 | 6.7 |

Niche overlap analyses indicated moderate overlap in geographic space (Schoener’s D= 0.498) and lower overlap in environmental space (D= 0.357). The niches were statistically different (p= 0.0099), and the observed overlap did not differ from random expectations (D: p= 0.178). Consistently, both species shared a central high-density niche core under warm and humid conditions, while showing species-specific niche extensions along environmental gradients (Fig. S3).

## Discussion

Our study provides one of the most comprehensive syntheses to date of the invasion history and potential distribution of the ambrosia beetles *E. fornicatus* and *E. perbrevis*. By integrating phylogeographic analyses, invasion chronology and ecological niche modeling, we show that these species have expanded far beyond their native range through multiple, largely independent introduction events. Despite their close phylogenetic relationship and similar native distributions, the two species exhibit contrasting invasion dynamics and occupy distinct, though partially overlapping climatic niches across native and non-native regions.

### Genetic structure and invasion history

Both species display a markedly structured genetic architecture, characterized by substantial differentiation among populations that suggest historical compartmentalization of genetic variation. The preponderance of monomorphic haplotypes with geographically restricted distributions points to two non-mutually exclusive processes: profound historical vicariance events, and recent colonization followed by accelerated genetic drift under isolation. The fixation of individual haplotypes across distant insular and continental territories constitutes compelling evidence of pronounced founder effects associated with long-distance, human-mediated dispersal. Such patterns are typical of invasive forest insects introduced via trade pathways and indicate limited contemporary gene flow among established populations (Estoup and Guillemaud 2010; Garnas et al. 2016; Urvois et al. 2023). Collectively, these results indicate that the global spread of both species has not resulted from a single global expansion, but rather from multiple introductions originating from geographically distinct source populations. Most successful invasions for both species can be traced directly to their native range, and while our data do not allow a formal test of bridgehead dynamics, the presence of genetically distinct lineages spreading among invaded regions is consistent with the possibility that some established populations may act as secondary sources of further introductions (Bertelsmeier and Keller 2018).

For *E. fornicatus*, the concentration of genetic diversity within China, Taiwan and Vietnam supports the hypothesis that this region represents the historical evolutionary reservoir of the species. Notably, the comparatively low genetic differentiation between China and Vietnam contrasts with the values observed toward most invasive populations, suggesting episodes of relatively recent gene exchange between these adjacent cores. We propose that this limited, yet discernible connectivity reflects historical permeability across a biogeographic transition zone that has remained conducive to biological dispersal under environmental change or anthropogenic disturbance.

The phylogenetic structure of *E. fornicatus* further indicates an invasion dynamic characterized by at least seven apparently independent introduction events into continental USA, Hawaii, South America, Central Europe, Spain, Turkey and South Africa, each associated with distinct haplotypes. Most of the introductions appear to originate directly from its native range (primary invasions), although there is also evidence of likely secondary spread between already invaded regions. Most introductions can be traced directly to Asian source populations (China, Taiwan and Vietnam), and notably, these introduction events appear to have occurred almost entirely within the last two decades. Although little information is available regarding the pest’s prevalence within its native range, this pattern may be explained by recent increases in population sizes and/or by the intensification of international trade in commodities that act as vectors for the species (i.e., the opening of new commercial pathways) (Hulme 2009; Essl et al. 2015).

In contrast, *E. perbrevis* exhibits higher overall genetic differentiation and more restricted gene flow, suggesting a more fragmented population structure. The high incidence of monomorphic records (67% of haplotypes occurring in single countries) supports scenarios involving either ancient vicariance events or recent colonization followed by rapid genetic drift in isolated populations. This pattern is particularly evident in Pacific Island populations, where single haplotypes have reached fixation (100% frequency), likely reflecting founder effects during sequential colonization events across this region. The substantial diversity observed in Thailand and Indonesia supports the hypothesis that these regions constitute the evolutionary core of this species complex (Smith et al. 2019). This elevated diversity could indicate: (i) an ancestral source population with high effective population size, (ii) a secondary contact zone where distinct lineages have converged, or (iii) a region experiencing ongoing gene flow from multiple sources. Papua New Guinea appears to represent a transitional region between two major lineages, a powerful model to study how ancestral polymorphisms are retained, partitioned, and influence adaptation within dynamic contact zones.

Outside the native range, *E. perbrevis* has established invasive lineages spanning a longer period of time than *E. fornicatus*. The most long-distance invasive populations of *E. perbrevis* appear to have originated from, or to be closely related to, the Thai lineage (e.g. haplotypes 12, 14 and 19) introduced into North America and Europe, and the Indonesian lineage (e.g. haplotypes 1, 2 and 23) introduced into Central America and Europe. In contrast, several island locations in Oceania, including American Samoa, Australia and Fiji, appear to result from independent introductions from currently unidentified source populations. Papua New Guinea and the Solomon Islands might represent a transitional region between two major lineages. Furthermore, the Indonesian lineage includes a sublineage related to Papuan populations, which shows a genetic affinity to the introduction in Vanuatu. Although *E. perbrevis* has been suggested to be native to northeastern Australia, with the southern records representing a natural range expansion (Li et al. 2024), the strong monomorphism and high differentiation observed in Australian populations relative to all other studied populations -including neighboring Pacific islands- are more consistent with an ancient, independent anthropogenic introduction followed by genetic drift. Similar patterns of pronounced population structuring and haplotype fixation following historical colonization events have been reported for other ambrosia beetle species (Gohli et al. 2016). Although limited sampling could potentially conceal cryptic diversity, this explanation is unlikely given the consistency of differentiation observed across all analyzed data.

Comparisons with other invasive ambrosia beetles are consistent with our findings. Studies on *Xylosandrus crassiusculus, X. germanus, X. compactus*, and *Euwallacea interjectus* indicate that introductions directly from the native range have been the dominant invasion pathway, while secondary spread has played a variable, though sometimes important, role (Dzurenko et al. 2021; Urvois et al. 2023; Lai et al. 2024). Collectively, these patterns underscore the predominance of Southeast Asia as a source region, while highlighting how multiple origins and occasional bridgehead dynamics can jointly shape global invasion histories.

### Climatic niches and potential distribution

Despite their ecological similarity and overlapping native distribution in SE Asia, *E. fornicatus* and *E. perbrevis* differ substantially in their climatic niches. The native range of *E. fornicatus* extends into more subtropical regions, while that of *E. perbrevis* is largely centered in tropical environments. Consistent with this, *E. fornicatus* has successfully invaded cooler and drier regions, suggesting broader climatic tolerance, potentially facilitated by human-mediated movement and conditions in greenhouses and urban areas (Ceriani-Nakamurakare et al. 2025). *Euwallacea perbrevis,* on the other hand, has established non-native populations in humid tropical regions, although isolated records in subtropical regions such as Florida and Panama, suggest that this species may also tolerate broader climatic conditions than previously assumed (Rabaglia et al. 2006; Gomez et al. 2018)(Rabaglia et al. 2006; Gomez et al. 2018)(Rabaglia et al. 2006; Gomez et al. 2018)(Rabaglia et al. 2006; Gomez et al. 2018).

The varying success of these species in establishing across different regions is likely linked to a combination of pathways of introduction (Meurisse et al. 2019) and differential climatic suitability. Our species distribution models support these differences, identifying temperature as key predictors of their global distributions. For instance, *E. fornicatus* showed high suitability across a wide thermal range (annual mean temperatures ranging from 11 to 26 °C), while *E. perbrevis* was more restricted to warmer (annual mean temperatures higher than 15 °C), showing lower tolerance to diurnal temperature variation. These patterns highlight the role of climate in limiting or facilitating the establishment of these beetles in new territories (Rassati et al. 2016; Lantschner et al. 2020). In line with this, our models predicted only limited overlap in climatic suitability (32%), indicating that coexistence is likely to be geographically constrained. The ranges of suitable temperatures for survival predicted by our models are consistent with laboratory studies. For *E. fornicatus*, the minimum and maximum temperatures required to complete development have been reported as 13 °C and 33 °C, respectively (Umeda and Paine 2019). In contrast, *E. perbrevis* has been shown to be unable to develop at minimum temperatures below 15 °C (Walgama and Zalucki 2007).

Based on our model predictions, the populations of *E. fornicatus* currently established in the United States of America (California) could find suitable habitat for further spread into the southeastern part of this country, eastern Mexico, and parts of Central America, although such an expansion would require crossing a substantial gap of unsuitable habitat. In South America, the species is predicted to encounter continuous suitable conditions allowing expansion into northern Argentina, Paraguay, and the eastern coast of Brazil. Likewise, populations in South Africa could potentially spread into temperate and subtropical regions of the northern part of the continent, while those established in Spain, Israel and Turkey may be able to colonize several areas across the Mediterranean basin. In Australia, suitable habitat is also predicted, particularly in western regions, although expansion would depend on the species’ ability to bridge current distribution gaps. For *E. perbrevis*, populations currently established in Central America may find favorable conditions for expansion into northern South America, including the Amazon basin and the Brazilian coast. Although the species has not yet been detected in Africa, our models indicate the existence of extensive areas of suitable habitat in central regions of the continent.

### Limitations and future directions

The invasion patterns inferred here are based on a single genetic marker (COI), which limits the resolution of introduction routes and the identification of source populations. More robust inferences would require the incorporation of additional genetic markers, such as nuclear loci, SNPs, or microsatellites, together with broader sampling across the native range to better capture the full spectrum of potential source populations. Similarly, predictions derived from species distribution models should be interpreted cautiously. The occurrence data used to calibrate these models may reflect constraints beyond climate that limit species occupancy, including biotic interactions such as competition, parasitism, and pathogen pressure, as well as land-use patterns (Beaumont et al. 2009). In addition, host availability and host-use patterns may further restrict the distribution of both species. Although they are highly polyphagous and capable of attacking and reproducing in a wide range of hosts, interspecific and regional variation in host preference may influence establishment success and subsequent spread.

Given the substantial genetic diversity documented in both species, together with growing evidence that genetic differentiation can be associated with functional divergence in traits such as host preference, management strategies that assume a single panmictic population model may be inadequate. While data on dispersal capacity and environmental tolerance remain limited, future studies integrating population genomics with ecological niche modeling and physiological studies will be essential to determine whether genetically distinct lineages differ in environmental requirements or invasion potential. In this context, some of the inherent limitations of correlative ecological niche models could be addressed in future studies by developing finer-scale mechanistic models that explicitly incorporate biological processes and species-specific traits, thereby improving predictions of establishment and spread under current and future environmental conditions (Morin and Thuiller 2009).

### Implications for management and biosecurity

The findings of this study enhance our understanding of the global invasion ecology of these two pests, the environmental factors shaping their distributions, and the regions where they may potentially invade in the future. Such insights can inform and strengthen surveillance and monitoring programs. It is noteworthy that, despite the stricter biosecurity measures implemented over the past two decades (e.g., ISPM 15), the long-distance movement of the studied species—particularly *E. fornicatus*—does not appear to have been significantly reduced. The main vectors reported for both species include untreated wooden articles, packaging materials, and plants for planting known to serve as hosts. In Europe, for example, trade of ornamental plants for botanical collections within the EU has been identified as the source of several outbreaks (Schuler et al. 2023). The repeated introduction of genetically distinct lineages mainly from its native range underscores the controls of these high-risk pathways.

Importantly, the contrasting outcomes of eradication efforts between climatically suitable and unsuitable regions emphasize the need for management strategies tailored to local environmental context. In climatically marginal areas, eradication efforts may be cost-effective and successful, while in climatically optimal regions, resources may be better allocated to containment and impact mitigation. An illustrative example is the management response to *E. fornicatus* in Perth, Australia, where intensive governmental eradication efforts were implemented over approximately four years, involving systematic removal and chipping of all infested trees. However, recent policy announcements indicate a strategic shift from eradication to containment, with management activities now focused on minimizing the risk of spread beyond the Perth metropolitan region (Australian Government 2025). The broader climatic tolerance of *E. fornicatus*, coupled with its rapid and ongoing global spread, suggests that this species may require more intensive and geographically extensive management efforts compared to the more climatically restricted *E. perbrevis*.

Overall, our results contribute to the understanding of ambrosia beetle invasion dynamics globally. The capacity for multiple, independent establishment events from diverse source populations appears to be a feature of successful invasive ambrosia beetles, potentially reflecting their association with international timber and/or ornamental trade and their ability to utilize diverse host plants. The phylogeographic evidence for climate-mediated establishment success also has implications for predicting future invasion risk under climate change scenarios. As global temperatures rise and precipitation patterns shift, the climatic suitability landscapes for both species might change dramatically. *Euwallacea fornicatus*, with its broader temperature tolerance, may experience range expansions into currently marginal temperate regions, while *E. perbrevis*, constrained by higher minimum temperature requirements, may face both range expansions in warming regions and contractions in areas experiencing reduced precipitation. These shifting suitability patterns will require adaptive management strategies that account for climate-driven changes in invasion risk.

## Conclusion

The contrasting invasion dynamics of *E. fornicatus* and *E. perbrevis* highlight the risks of treating cryptic species complexes as ecologically equivalent. Despite their close phylogenetic relationship, these ambrosia beetles differ markedly in their invasion trajectories, climatic tolerances, and rates of spread, underscoring the need for taxonomically resolved and species-specific management strategies. Their divergent invasion patterns emphasise the combined role of climate and human activity in shaping their global spread, likely reflecting distinct trade dynamics for both species, often aligned with regions of climatic suitability. *Euwallacea fornicatus* exhibits broad climatic tolerance, enabling its successful establishment in both temperate and subtropical regions. Conversely, *E. perbrevis* remains more restricted to warm, humid climates, though some records in subtropical zones suggest flexibility, likely through exploitation of microclimates. Molecular phylogenetic analysis underscores the intricacies of the anthropogenic and natural dispersal of ambrosia beetles, where divergent lineages are being established on a global scale from diverse source populations.

## Acknowledgments

We would like to thank all the researchers who generously shared novel molecular information. This work is part of a MoU and MTA between the University of Florida (United States) and the School of Agriculture of the University of Buenos Aires (Argentina).

## Author Contribution Statement

MVL, ECN, AJJ and DFG conceptualized and designed the study. AIC and SMS participated in sample collection and obtained the sequence data. MVL conducted the species distribution models. ECN conducted the population genetic analysis. MVL and ECN analyzed the data. MVL, ECN and DFG wrote the first draft. All authors reviewed and edited the manuscript. All authors gave final approval for publication and agreed to be held accountable for the work performed therein.

## Data Availability

All data supporting the findings of this study are available in the Supplementary Information associated with this article.

## Supplementary Information

**Figure S1** Geographic distribution and relative frequencies of mitochondrial haplotypes from *Euwallacea fornicatus* and *Euwallacea perbrevis*. Pie charts represent the relative frequency of haplotypes within each sampled country. Country codes follow the ISO 3166-1 alpha-3 standard. For the United States (USA), samples include both continental and insular territories (e.g., Hawaii). Numbers beneath each pie chart indicate haplotype identities present in that population (e.g., “1” indicates haplotype H1; “8-10” indicates haplotypes H8, H9 and H10)

**Figure S2** Response curves showing the relationship between the five bioclimatic variables included in the distribution models of *E. fornicatus* and *E. perbrevis* (see Table 1 for details) and the predicted suitability from the TSS-weighted ensemble model (GBM, GLM, MaxEnt, and RF). The curves illustrate how predicted suitability changes as each environmental variable varies, while all other variables are held constant at their median values. Solid black lines represent the mean prediction across all models and replicates, and grey shaded areas indicate ±1 standard deviation around the mean.

Figure S3 Climatic niche overlap between *Euwallacea fornicatus* (left) and *Euwallacea perbrevis* (right) in the multivariate environmental space defined by the first two principal components (PC1 and PC2) of a PCA of global bioclimatic conditions. The blue gradient represents the kernel density of occurrence records for the focal species (darker= higher density of occurrences). Green areas indicate the portion of the focal species’ niche that is not occupied by the other species (unique niche space). Red areas indicate the portion of the other species’ niche that is not occupied by the focal species (niche of the compared species not shared with the focal species). Blue-purple areas indicate shared niche space (overlap). The dark red contour line delimits the extent of the globally available environmental space sampled as background.

**Table S1** Occurrence records of *Euwallacea fornicatus* and *Euwallacea perbrevis* used in this study. We include country, administrative division, locality, geographic coordinates, year of collection, native status, data source (see full bibliographic list), identification method (morphological or genetic), and corresponding GenBank accession numbers when available. Records include both published and unpublished data from native and non-native ranges

**Table S2** Haplotype metadata for *Euwallacea fornicatus* based on mitochondrial COI sequences. For each haplotype, the table reports the haplotype identifier, its frequency across countries, and the corresponding GenBank accession number followed by the country of origin. Country codes: ARG: Argentina, AUS: Australia, BRA: Brazil, CHN: China, DEU: Germany, ESP: Spain, ISR: Israel, ITA: Italy, JPN: Japan, NLD: Netherlands, POL: Poland, THA: Thailand, TUR: Turkey, TWN: Taiwan, URY: Uruguay, USA: United States of America, VNM: Vietnam, ZAF: South Africa

**Table S3** Pairwise genetic differentiation among *Euwallacea fornicatus* populations based on mitochondrial COI sequences. Estimates include haplotype diversity (Hs), nucleotide diversity within populations (Ks), mean number of nucleotide differences between populations (Kxy), and multiple measures of genetic differentiation and structure (Gst, ΔSt, ΓSt, Nst, Fst), as well as absolute (Dxy) and net nucleotide divergence (Da). Country codes: ARG: Argentina, AUS: Australia, BRA: Brazil, CHN: China, DEU: Germany, ESP: Spain, ISR: Israel, ITA: Italy, JPN: Japan, NLD: Netherlands, POL: Poland, THA: Thailand, TUR: Turkey, TWN: Taiwan, URY: Uruguay, USA: United States of America, USAi: United States of America (Hawaii), VNM: Vietnam

**Table S4** Haplotype metadata for *Euwallacea perbrevis* based on mitochondrial COI sequences. For each haplotype, the table reports the haplotype identifier, its frequency across countries, and the corresponding GenBank accession number followed by the country of origin. Country codes: AUS: Australia, ASM: American Samoa, CHN: China, CRI: Costa Rica, FJI: Fiji, IDN: Indonesia, LKA: Sri Lanka, NLD: Netherlands, PAN: Panama, PNG: Papua New Guinea, SLB: Solomon Islands, THA: Thailand, USA: United States of America, VUT: Vanuatu

**Table S5** Pairwise genetic differentiation among *Euwallacea perbrevis* populations based on mitochondrial COI sequences. Estimates include haplotype diversity (Hs), nucleotide diversity within populations (Ks), mean number of nucleotide differences between populations (Kxy), and multiple measures of genetic differentiation and structure (Gst, ΔSt, ΓSt, Nst, Fst), as well as absolute (Dxy) and net nucleotide divergence (Da). Country codes: AUS: Australia, ASM: American Samoa, CHN: China, CRI: Costa Rica, FJI: Fiji, IDN: Indonesia, LKA: Sri Lanka, NLD: Netherlands, PAN: Panama, PNG: Papua New Guinea, SLB: Solomon Islands, THA: Thailand, USA: United States of America, VUT: Vanuatu

**Table S6** Predictive performance of individual SDMs for *Euwallacea fornicatus* and *E. perbrevis*, evaluated through 5-fold cross-validation with 5 repetitions (n= 27 replicates per algorithm). Values represent mean ± standard deviation of calibration scores across replicates. GLM: Generalized Linear Models, RF: Random Forest, GBM: Gradient Boosting Machines, and MaxEnt: Maximum Entropy.

**Table S7**. Performance of MaxEnt models for *Euwallacea fornicatus* and *E. perbrevis* across combinations of regularization multipliers (RM= 0.5–5) and feature classes (FC: L= linear, Q= quadratic, H= hinge, P= product). Model performance is reported as mean AUC (AUC_avg ± AUC_sd), along with minimum (AUC_min) and maximum (AUC_max) values across cross-validation replicates.

**Table S8.** Relative contribution of each algorithm to the TSS-weighted ensemble model for each species. GLM: Generalized Linear Models, RF: Random Forest, GBM: Gradient Boosting Machines, and MaxEnt: Maximum Entropy.

